## Supplementary Figures for "Evolutionary plasticity and functional repurposing of the essential metabolic enzyme MoeA"

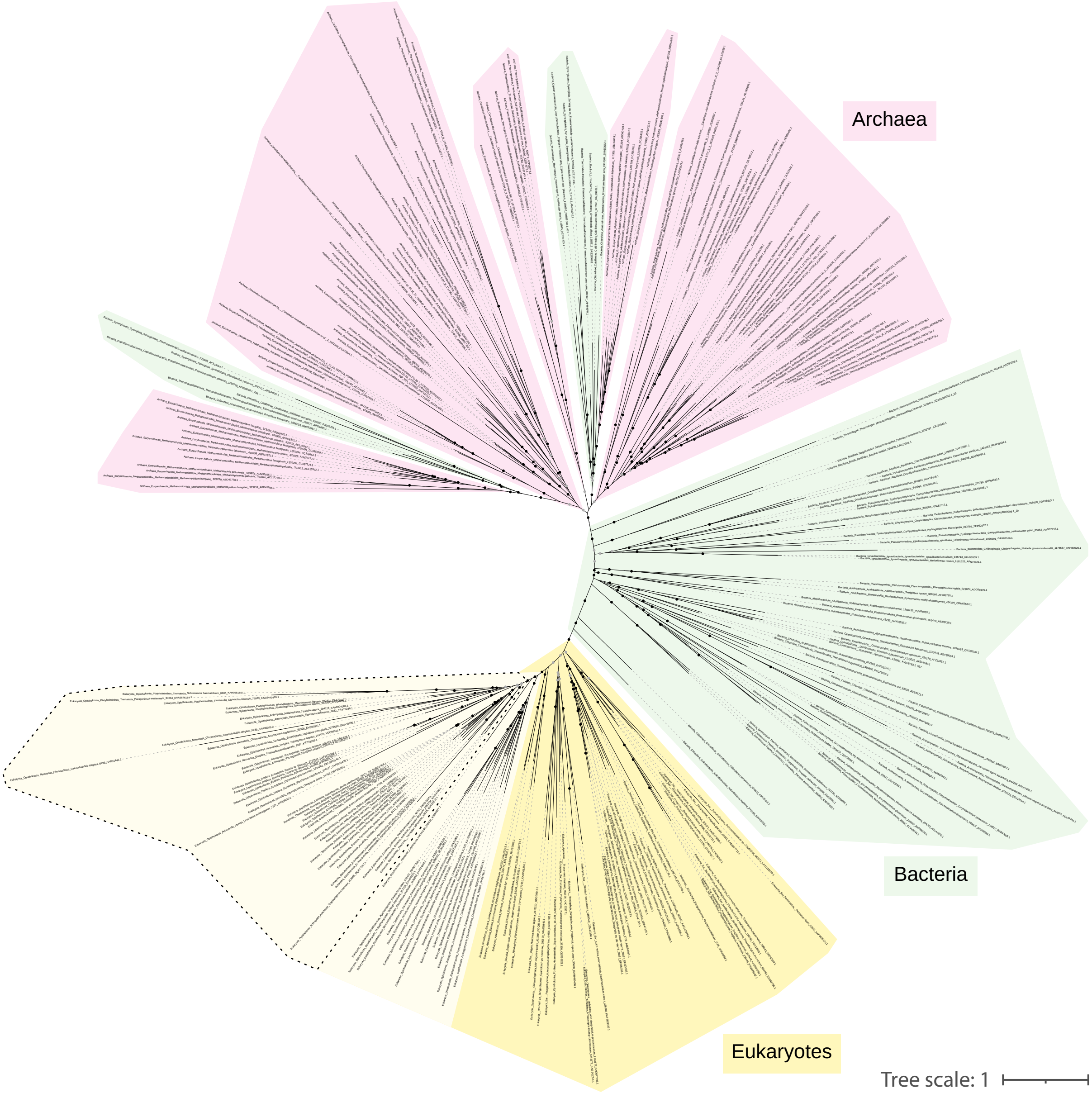


**Supplementary Figure 1**

Maximum-likelihood phylogeny of MoeA/Gephyrin in Eukaryotes, Bacteria and Archaea. Black dots indicate UFB > 90, gray dots indicate 80 < UFB <= 90 and branches without dots indicated UFB <= 80. The scale bar represents the average number of substitutions per site.

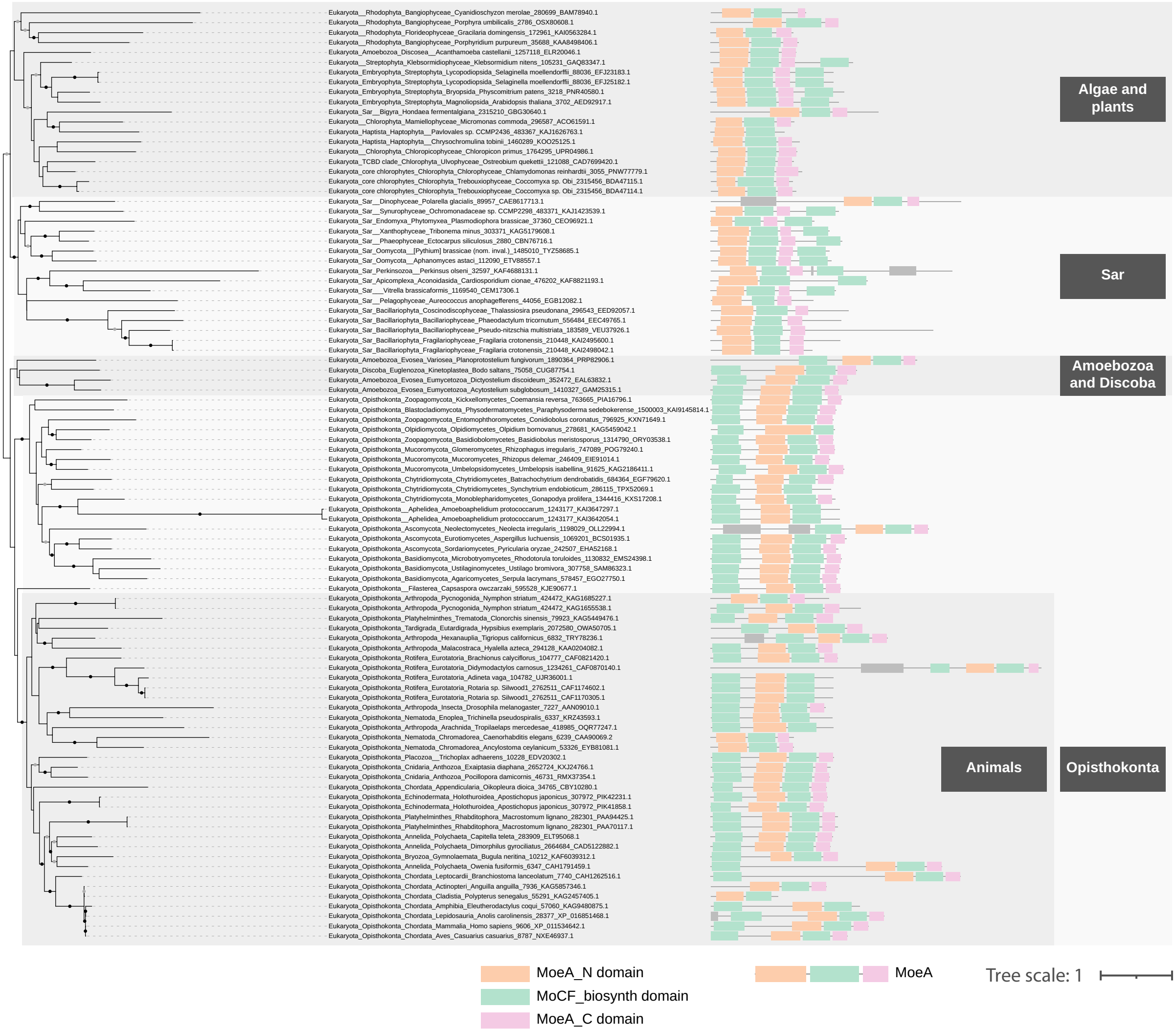


### Supplementary Figure 2

Domain organization of MoeA/Gephyrin in a maximum-likelihood phylogeny of MoeA/Gephyrin in Eukaryotes. Domains indicated in gray correspond to domains different to MoeA or MogA. Higher taxonomic ranks are indicated in black boxes. Black dots indicate UFB > 90, gray dots indicate 80 < UFB <= 90 and branches without dots indicated UFB <= 80. The scale bar represents the average number of substitutions per site.

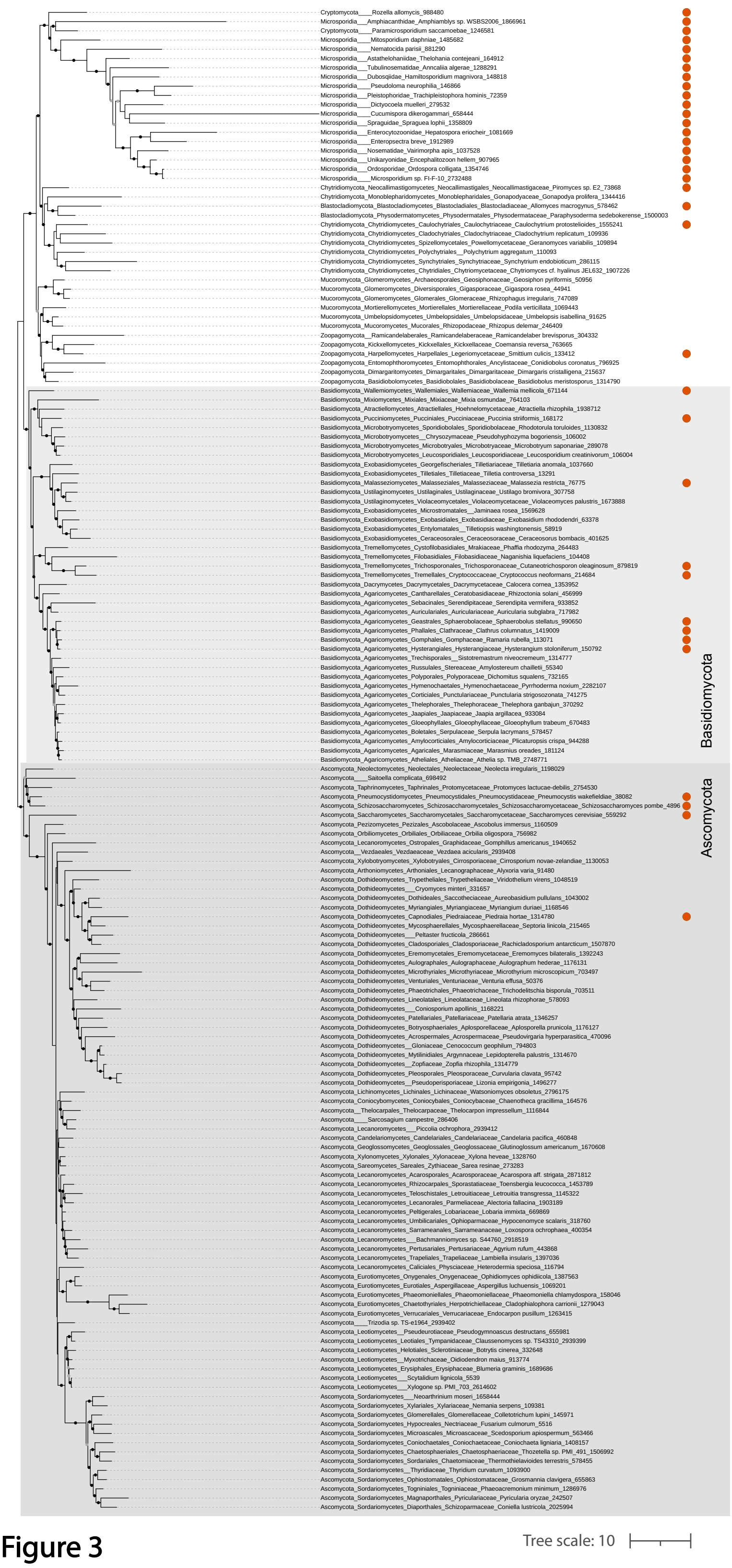


### Supplementary Figure 3

Phyletic pattern of the absence of MoeA in kingdom Fungi. The light gray box indicates the clade corresponding to phylum Basidiomycota, and the dark gray box indicates phylum Ascomycota. For the detailed information, see Supplementary Table 1. Black dots indicate UFB > 90, gray dots indicate 80 < UFB <= 90 and branches without dots indicated UFB <= 80. The scale bar represents the average number of substitutions per site.

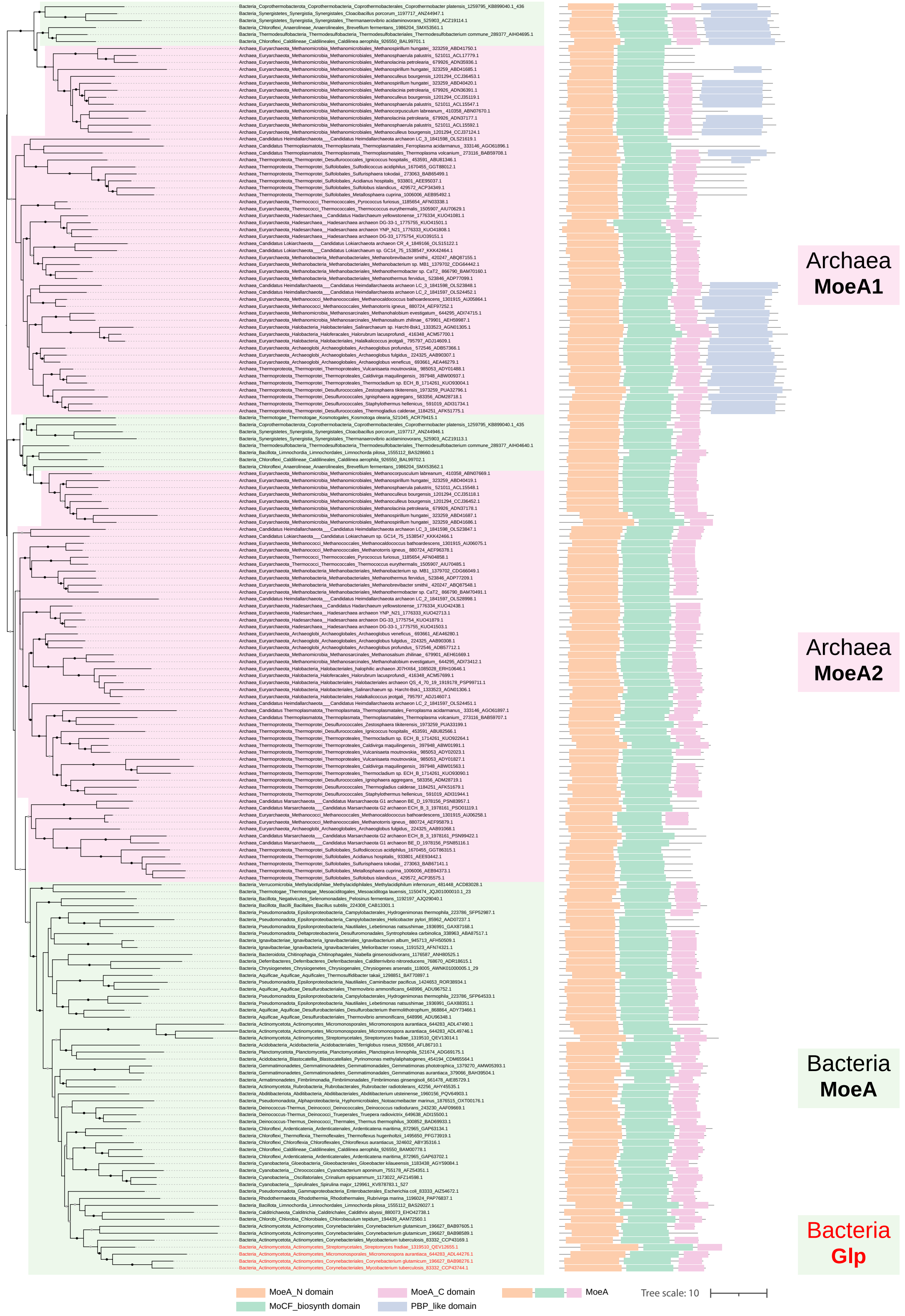


### Supplementary Figure 4

Maximum-likelihood phylogeny of MoeA in Archaea and Bacteria with a schematic representation of the domain organization of MoeA. Bacteria phyla are indicated in green, and Archaea phyla are indicated in pink. Black dots indicate UFB > 90, gray dots indicate 80 < UFB <= 90 and branches without dots indicated UFB <= 80. The scale bar represents the average number of substitutions per site.

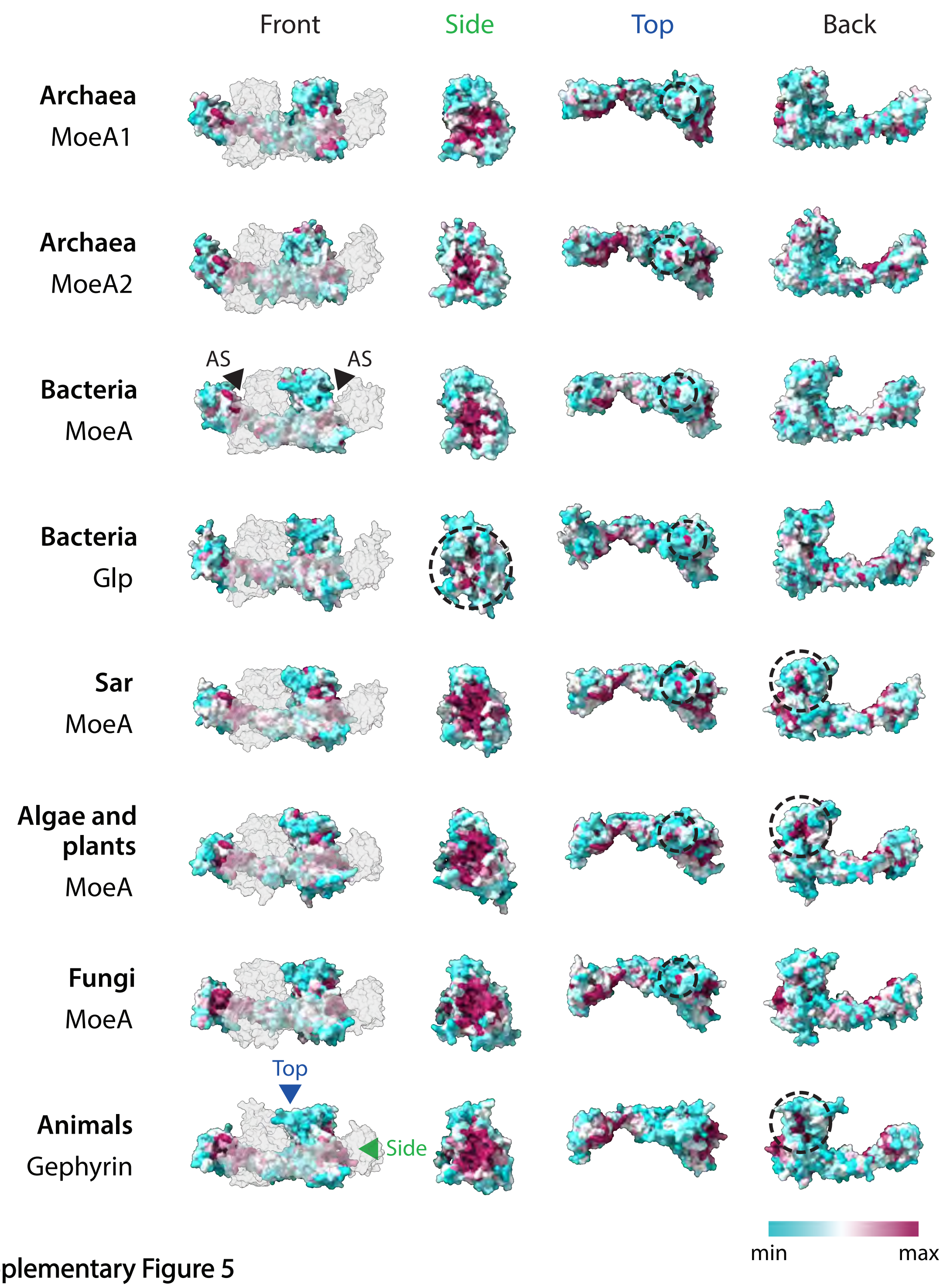


**Supplementary Figure 5**

Sequence conservation of MoeA mapped on a representative structure of each group indicated on the left. In the front view, the second subunit of the monomer is indicated in gray. Arrows in Gephyrin point to the direction of the Side and Top view of the structure. The dashed line in Glp indicates a decreased sequence conservation in the putative active site. The dashed line in Gephyrin indicates a high sequence conservation in the neuroreceptor binding domain. Note that this region is also well conserved in Sar, algae and plants. Dashed lines in the top view indicate a highly conserved Gly residue in all studied groups except animals. When present, PBP and MogA domains were hidden to facilitate the visualization of MoeA. AS: Active site.
